# Acyl Carrier Protein is essential for MukBEF action in *Escherichia coli* chromosome organization-segregation

**DOI:** 10.1101/2021.04.12.439405

**Authors:** Josh P. Prince, Jani R. Bolla, Gemma L. M. Fisher, Jarno Mäkelä, Carol V. Robinson, Lidia K. Arciszewska, David J. Sherratt

## Abstract

Structural Maintenance of Chromosomes (SMC) complexes contribute ubiquitously to chromosome organization-segregation. SMC proteins have a conserved architecture, with a dimerization hinge and an ATPase head domain separated by a long antiparallel intramolecular coiled-coil. Dimeric SMC proteins interact with essential accessory proteins, kleisins that bridge the two subunits of an SMC dimer, and HAWK/KITE accessory proteins that interact with kleisins. The ATPase activity of the *Escherichia coli* SMC protein, MukB, is essential for *in vivo* function and is regulated by interactions with its dimeric kleisin, MukF, and KITE, MukE. Here we demonstrate that, in addition, MukB interacts with Acyl Carrier Protein (AcpP) that has essential functions in fatty acid synthesis. We characterize the AcpP interaction site at the joint of the MukB coiled-coil and show that the interaction is essential for MukB ATPase and for MukBEF function *in vivo*. Therefore, AcpP is an essential co-factor for MukBEF action in chromosome organization-segregation.

## Introduction

In *Escherichia coli*, the SMC complex, MukBEF, is composed of three essential proteins, the SMC protein MukB, the kleisin, MukF and the KITE protein, MukE^1-3^. Although divergent in primary sequence from other SMC proteins, MukB shares common ancestral and architectural features including an ABC-like ATPase head domain, a ∼50 nm long antiparallel coiled-coil and a dimerization hinge domain (Fig. 1a). In addition, MukB retains two highly conserved discontinuities within the coiled-coils. The first, the ‘joint’, located ∼100 amino acids from the head domain, is highly conserved between SMC complexes, and has been suggested to aid flexibility for head engagement during ATP hydrolysis cycles^4-8^. The other, roughly half-way along the coiled-coils, the ‘elbow’, enables the protein to fold upon itself bringing the hinge domain in close proximity to one of the two ATPase heads, though the functional implications of this are unclear^5,9-11^. As with other SMC proteins, MukB dimers interact with their klesin, MukF, through two distinct interaction sites; one in the ‘neck’ region of the coiled-coils, located between the head and the joint of one monomer, and the ‘cap’ region of the partner ATPase head (Fig. 1a)^2,12,13^. Unusually among kleisins, MukF dimerizes through an additional N-terminal dimeric winged-helix domain (WHD). This enables the joining of two dimeric MukBEF complexes into dimer of dimer (DoD) complexes that are essential for *in vivo* MukBEF function^3,13,14^. MukE dimers interact with MukF; thus the complete MukBEF complex has a 4:4:2 B:E:F stoichiometry^3,14^. MukB ATP hydrolysis results from the engagement of two head domains that create two shared ATP binding sites. MukB alone has minimal ATPase activity but is activated in the presence of MukF and further modulated by the interactions with MukE and DNA^12^. ATPase activity is essential for *in vivo* function, as mutant MukB proteins deficient in ATP hydrolysis (MukB^E1407Q^, hereafter referred to as MukB^EQ^) or ATP binding (MukB^D1407A^, hereafter referred to as MukB^DA^) display *ΔmukB* phenotypes^3,15,16^.

**Figure 1.**
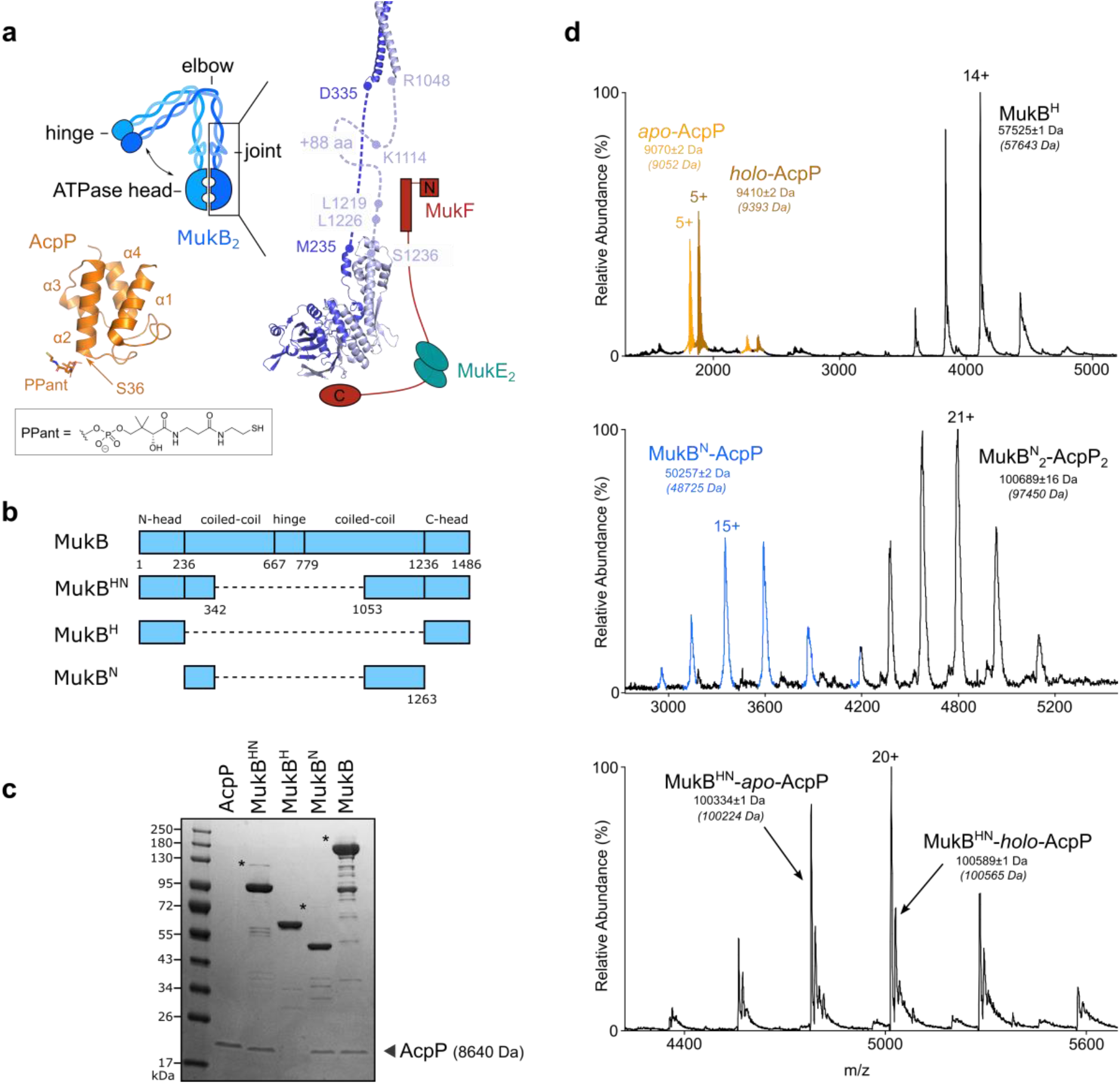
Specific binding of AcpP to the neck region of MukB. (**a**) Schematic of MukBEF in the elbow bent configuration (left); structure of *E. coli* MukB^HN^ (right) using crystal structure of the elbow (PDB 6H2X)^9^, and a homology model based on *H. ducreyi* MukB^H^ structure (PBD 3EUK)^2^. The coiled-coil and joint are modelled and MukEF are shown in cartoon form (note that the C- and N-terminal domains of a given MukF monomer normally contact different MukB monomers). Structures of AcpP-PPant are also shown (bottom, left, PBD 3NY7)^48^. (**b**) Schematic of MukB truncations (**c**) SDS-PAGE analysis of AcpP co-purification with MukB truncations. Putative disulfide linked MukB-AcpP species are indicated with an asterisk. Note that AcpP (MW 8640 Da) runs with an apparent MW of ∼18000 Da on SDS-PAGE. (**d**) nMS analysis of AcpP-MukB truncation interactions. Top, MukB^H^ with the addition of recombinant AcpP (mixed population of *apo* and *holo* species), middle, MukB^N^ with copurified AcpP and bottom, MukB^HN^ with copurified AcpP. Theoretical masses in parentheses.

Acyl Carrier Protein (AcpP) has been repeatedly reported to co-purify with MukB^14,17-19^. Since AcpP is a highly abundant *E. coli* protein (1-36 × 10^4^ molecules/cell; >100 times excess over endogenous MukB)^3,20-22^, it was not clear from early reports whether this reflected a specific interaction or a fortuitous association. AcpP is an essential hub protein that through a covalent interaction with its phosphopantetheine (PPant) arm, shuttles intermediates along the fatty acid biosynthesis pathway by a series of acyl transfer reactions (Fig. 1a) (reviewed in^23^). In addition, AcpP has been shown to interact with other unrelated protein partners including SpoT, IscS and SecA^24-26^. Searches for binding partners of AcpP have also indicated an interaction with MukB, although any functional significance to this interaction was not explored^24,26,27^.

Here, we identify the AcpP binding site on MukB and analyze the functional consequences of this interaction *in vitro* and *in vivo*. We show that the interaction of AcpP with a conserved region in the MukB joint within the coiled-coils is essential for MukB ATPase activity. The binding of AcpP to MukB inhibits higher order intermolecular interactions *in vitro* between MukB and MukB^HN^ (MukB Head-Neck, containing the MukB ATPase head plus ∼30% of the adjacent coiled-coils). Mutations within the AcpP binding site reduce AcpP association and thus impair MukB ATPase activity. Importantly, these mutations result in an altered pattern of MukBEF complex localization within cells, including an increased association with the replication termination region (*ter)*, consistent with the impaired ATPase function. The data lead to the conclusion that AcpP is an essential partner in MukBEF function.

## Results

### AcpP interacts with the MukB coiled-coils

The nature and function of the interaction between AcpP and MukB has been unclear, despite numerous reports describing an interaction^14,17,19,24,27,28^. We therefore set out to determine whether the interaction between AcpP and MukB is specific and to identify any interaction site on MukB. Wild type (WT) MukB and three truncated variants were purified and tested for the presence of associated AcpP using SDS-PAGE (Fig. 1b, c). Because previous work had shown that truncated MukB hinge mutants did not co-purify with AcpP^15,29,30^, we focused on variants containing the ATPase head and head proximal regions. AcpP co-purified with MukB^HN^ (MukB Head-Neck) consisting of the ATPase head and first ∼30% of head-proximal coiled-coils, but not with MukB^H^ (MukB Head), consisting of just the ATPase head domain. Even with the addition of recombinant AcpP, no MukB^H^-AcpP binary complexes were detected. Consistent with these observations, AcpP co-purified with MukB^N^ (MukB Neck) consisting of just the head-proximal coiled-coils (Fig. 1c). Analysis of samples containing AcpP and MukB^N^ or MukB^HN^ using native Mass Spectrometry (nMS) revealed AcpP interacts with MukB with a 1:1 monomer-monomer stoichiometry (Fig. 1d), supporting data previously reported for WT MukB^14^. In addition, complexes with a mass corresponding to MukB^N^_2_-AcpP_2_ were also identified, likely arising through interactions between the coiled-coils. No such dimers were detected in MukB^HN^-AcpP samples.

To identify the MukB-AcpP interface, we utilized *in vitro* chemical cross-link mass spectrometry (XL-MS). Treatment of MukB^HN^ with BS^3^ cross-linker in the absence of AcpP, generated a mixture of inter- and intra-molecular cross-links (Supplementary Fig. 1a). In the presence of AcpP, despite the lack of detected MukB^HN^-AcpP cross-links, we noted the disappearance of three substantial species; their analysis showed that the presence of AcpP inhibited the formation of four cross-linked species involving residue K1125 (Supplementary Fig. 1b). This residue is located within the C-terminal helix in the coiled-coil proximal to the ATPase head domain and is present in both MukB^N^ and MukB^HN^ truncations (Fig. 1a, b and Supplementary Fig. 1b). Crystal structures of the MukB elbow and ATPase head indicates the C-terminal helix in this region includes an additional ∼80 residues compared to the N-terminal helix and likely forms a conserved ‘joint’ motif, also evident in cross-linking experiments (Fig. 1a)^4,5,6,11^. Sequence alignment of MukB proteins around K1125, indicates a high conservation of this and basic residues K1114 and R1122 (Supplementary Fig. 1c).

Other characterized AcpP-partner protein interfaces often involve electrostatic interactions centered on the α2 helix^31,32^ (Fig. 1a). This also seems to be true for the MukB-AcpP interface, as substitutions in the α2 helix of AcpP abolished its co-purification with MukB^24,28^. Hence, we reasoned these highly conserved basic residues in MukB might well comprise at least part of the MukB-AcpP interface. Accordingly, we sequentially mutated residues K1114 - K1125 to glutamic acid in an attempt to perturb the AcpP-MukB interface. In addition, we constructed a double and triple charge reversed MukB mutant, MukB^KK^ (containing the K1114E and K1125E mutations) and MukB^KRK^ (containing K1114E, R1122E and K1125E mutations). We observed a reduction in the levels of co-purified AcpP in MukB^K1114E^, MukB^W1117E^ and MukB^C1118E^ samples, as judged by SDS-PAGE, confirming the importance of these residues to the MukB-AcpP interface (Supplementary Fig. 2c and 2d). We also observed a loss of AcpP co-purification in the MukB^KK^ and MukB^KRK^ samples. Together, these results provide strong evidence for a specific AcpP binding site located at the joint, within the MukB coiled-coils.

### AcpP stimulates MukB ATPase activity *in vitro*

To characterize the functional significance of the MukB-AcpP interaction, AcpP was depleted from WT MukB during heparin purification using an extended salt gradient, where AcpP-depleted MukB eluted as a second peak with a higher retention time (Supplementary Fig. 2a and 2b). We then sought to identify any effects of removing AcpP on the ATPase activity of MukB. No detectable ATP hydrolysis was observed for the AcpP-depleted MukB sample and only minimal activity was seen as a result of MukB activation by MukF (2.0 ± 1.4 ATP molecules/MukB_2_/min; Fig. 2a and 2b). Remarkably, addition of recombinant AcpP restored ATPase activity (3.3 ± 0.6 and 29.0 ± 1.6 ATP molecules/MukB_2_/min respectively), to a level comparable with MukB and MukBF co-purified with AcpP (3.7 ± 0.2 and 27.2 ± 1.2 ATP molecules/MukB_2_/min respectively) and similar to that reported previously (where the samples will have contained co-purified AcpP)^12^. Consistent with this, addition of MukE to AcpP containing MukBF samples modestly inhibited MukF activation (Fig. 2a and 2b), as reported previously^12^. In these experiments, recombinant AcpP was a mixed population of *apo-* and *holo-*AcpP (lacking or containing the PPant prosthetic group, respectively). The relative contributions of these forms are explored later.

**Figure 2.**
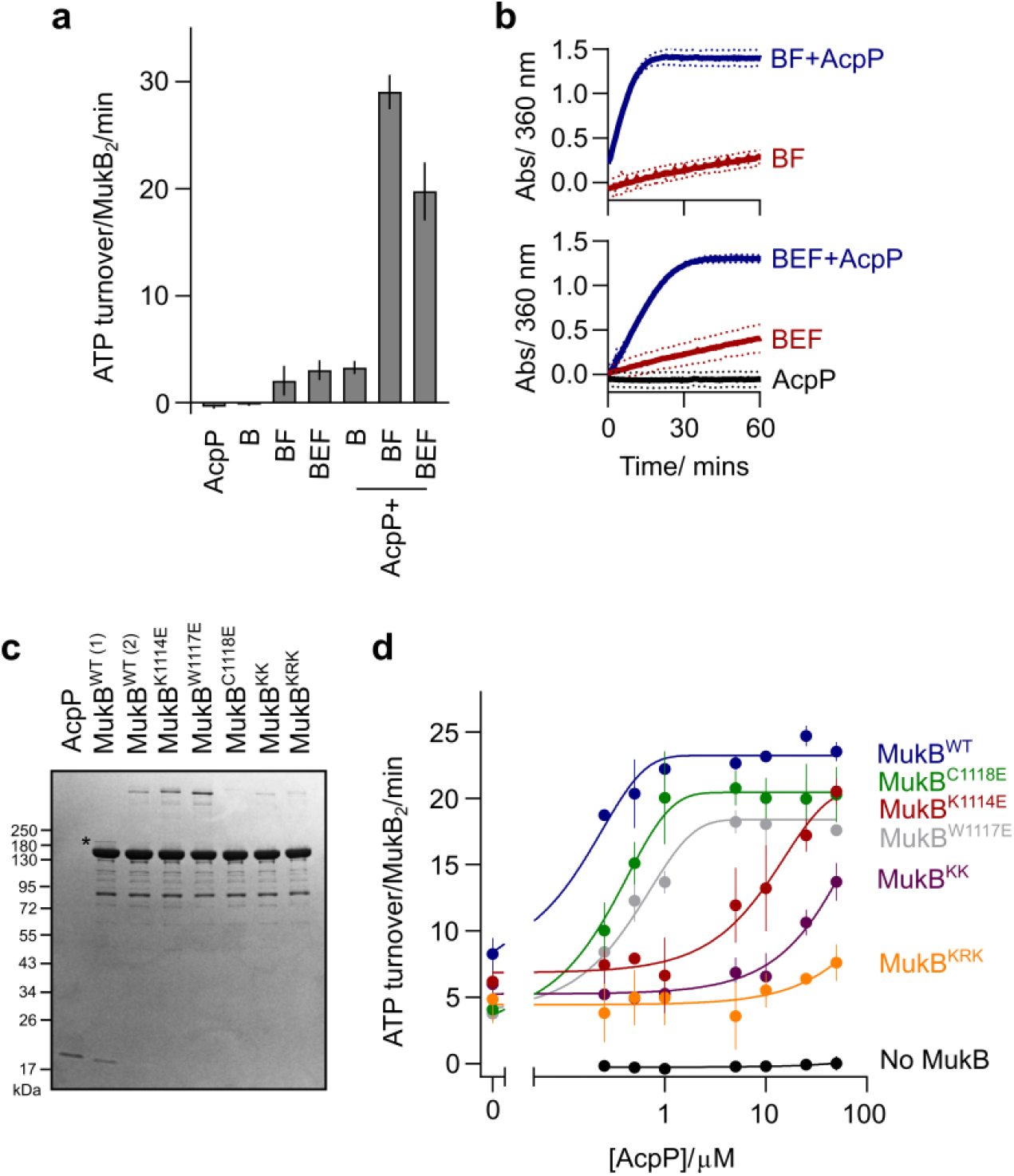
MukB ATPase activity requires interaction with AcpP. (**a**) Initial rate ATPase activity measurements of MukB in the presence and absence of AcpP (±SD from 3 technical repeats). (**b**) Absorbance data showing the measured activity of MukB over a time course of 60 min. (**c**) SDS-PAGE analysis of purified MukB variants highlighting the absence of copurified AcpP. MukB WT (1) and (2) refer to protein isolated from the heparin column from peaks 1 and 2 (Supplementary Fig. 1). Putative disulfide linked MukB-AcpP species are indicated with an asterisk. (**d**) Initial rate ATPase activity measurements of MukB proteins in response to increasing concentrations of AcpP (±SD from 3 technical repeats). MukF and MukE were included at a constant concentration in all samples.

As the presence of both AcpP and MukF was required for the activation of MukB ATPase activity, we considered the possibility that the binding of AcpP to MukB was a prerequisite for the interaction with MukF. Note that residues K1114-K1125 in the vicinity of the AcpP binding site are ∼100 residues N-terminal of L1219 and L1226, which have been implicated in MukF N-terminal binding^12^. We utilized nMS and blue native gel electrophoresis (BN-PAGE) to identify the formation of MukBEF complexes *in vitro* and to assess if AcpP influences MukEF binding. nMS analysis of mixtures of MukB, E, F and AcpP identified complexes consistent with a MukB_2_E_4_F_2_ stoichiometry with one or two AcpP molecules bound. In addition, complexes with masses corresponding to MukB_4_E_4_F_2_ and three or four molecules of bound AcpP were also observed (Fig. 3a). These dimer of dimer (DoD) complexes arise when a MukF dimer binds two separate MukB dimers (Fig. 3b). Consistent with this, increasing the concentration of MukB led to a higher proportion of DoD complexes. Complementary BN-PAGE experiments with a momomeric MukF derivative^12^, confirmed that DoD complexes depend on MukF dimerization (Fig. 3d). Furthermore, the formation of dimer or DoD complexes was independent of AcpP, but dependent on the MukB concentration (Fig. 3c); thereby demonstrating that AcpP binding to MukB is not required for the interaction with MukEF. These experiments also show that the formation of MukBEF DoD complexes requires neither bound nucleotide, nor head engagement (see Discussion).

**Figure 3.**
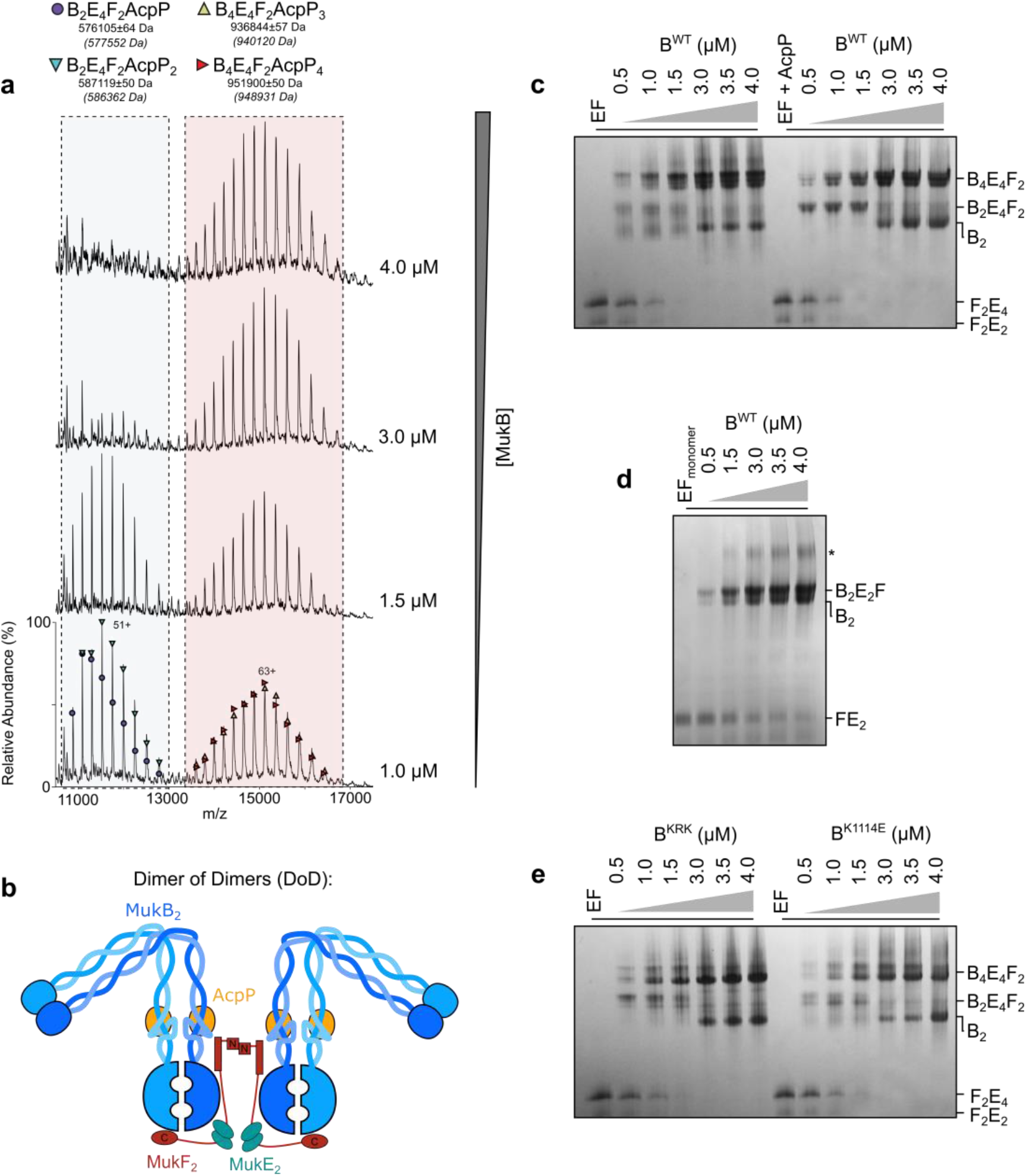
MukBEF forms DoD complexes independent of AcpP binding. (**a**) nMS analysis of MukBEF-AcpP complexes at various concentrations of MukB. (**b**) Schematic of MukBEF DoD complexes, approximate position of the AcpP binding site is indicated. (**c**-**e**) BN-PAGE analysis of complex formation in MukBEF-AcpP showing; (**c**) DoD complex formation is not dependent on the presence of AcpP, (**d**) higher order, DoD, complexes require the presence of dimeric MukF and (**e**) MukB^KRK^ and MukB^K1114E^ still form DoD complexes. Note co-purification of endogenous MukF with recombinant MukB led to the formation of MukB_4_E_4_F_2_ complexes in samples containing monomeric F, indicated with an asterisk.

MukB^K1114E^ and MukB^KRK^ mutant proteins, deficient in AcpP binding, were also able to form complexes with MukEF (Fig. 3e). Additionally, MukEF formed complexes with MukB^HN^ variant proteins (K1114E and C1118E) (Supplementary Fig. 3d). These data, taken together with the results of *in vivo* analysis showing formation of chromosome-associated MukB^KRK^EF foci (later), demonstrate that the AcpP interaction with MukB is not a prerequisite for MukF binding. Instead AcpP acts independently to regulate MukBEF activity in a way that remains to be mechanistically determined.

### Mutagenesis of the MukB joint region impairs AcpP-activated ATPase

To further analyze the requirement of AcpP binding for MukB ATPase activity, we analyzed the mutant proteins that failed to co-purify with AcpP (MukB^K1114E^, MukB^W1117E^, MukB^C1118E^, MukB^KK^ and MukB^KRK^) (Fig. 2c and Supplementary Fig. 2c and 2d). All five mutant proteins showed low ATPase activity in the presence of MukEF, in contrast to the mutants that co-purified with AcpP, which exhibited levels consistent with the amount of AcpP present within the sample (compare Supplementary Fig. 2d and 2e). The mutant proteins that lacked co-purified AcpP were then tested to see if the addition of recombinant AcpP stimulated their ATPase activity. AcpP-depleted WT MukB regained maximal ATPase activity after the addition of a 2-fold molar excess of AcpP (Fig. 2d). MukB^W1117E^ and MukB^C1118E^ both regained maximal ATPase activity with a 2-10 fold molar excess of AcpP, suggesting that these substitutions had only a modest impact on the MukB-AcpP interface, despite the conservation of these residues in MukB proteins (Supplementary Fig. 1c). The charge reversal mutants, MukB^K1114E^, MukB^KK^ and MukB^KRK^, showed a sequential reduction in the ability of AcpP to stimulate activity. At 100 times AcpP excess (but approaching the cellular concentration) the activity of MukB^KRK^ was only 7.6 ± 1.4 ATP molecules/MukB_2_/min (∼24% of the WT MukBF level in the presence of AcpP) (Fig. 2d). These data support the conclusion that AcpP binding to MukB is essential for *in vitro* ATPase activity.

### AcpP binding to the MukB joint inhibits intermolecular interactions

The initial assignment of the AcpP binding site on MukB was inferred from XL-MS experiments that showed that AcpP binding perturbs the formation of a BS^3^-induced intermolecular cross-link between two K1125 MukB^HN^ residues in the joint (Supplementary Fig. 1a and 1b). Discrete intermolecular MukB^HN^ complexes that disappeared in the presence of AcpP, were also observed in BN-PAGE of MukB^HN^EF mixtures (Supplementary Fig. 3a). We propose that these latter complexes have a stoichiometry of MukB^HN^_4_E_8_F_4_ and arise from the dimerization of MukB^HN^_2_E_4_F_2_ complexes through coiled-coil interactions in the region of the joint where the K1125 residues were cross-linked by BS^3^ (Supplementary Fig. 3e). Addition of AMPPNP to mixtures containing MukB^HN^, MukEF and AcpP, led to head engagement between AcpP-containing MukB^HN^_2_E_4_F_2_ complexes, resulting in MukB^HN^_4_E_4_F_2_AcpP_3/4_ complexes, equivalent to DoD complexes for WT MukB^14^ (Supplementary Fig. 3b and 3c). In the absence of AcpP, AMPPNP-induced head engagement led to the formation of presumptive MukB^HN^_8_E_8_F_4_ higher order complexes, arising through the same intermolecular coiled-coil interactions, as in the absence of AMPPNP (Supplementary Fig. 3c).

The single substitution mutant proteins (K1114E and C1118E) failed to produce detectable AMPPNP-dependent higher order complexes, irrespective of the presence of AcpP (Supplementary Fig. 3d), suggesting that the glutamate substitution in these proteins is sufficient to disrupt their intermolecular interaction. Consistent with AcpP perturbing intermolecular coiled-coil interactions between joint regions, we observed that higher order bands, formed through a presumptive disulfide interaction between two C1118 residues, were also inhibited by AcpP (Fig. 2c and Supplementary Fig. 2b). Any functional significance of the intermolecular interactions between the coiled-coil joint regions observed here and their inhibition by AcpP remains to be determined, as does understanding whether the inhibition by AcpP is a consequence of a steric constraint, or by AcpP inducing a conformational change in the MukB coiled-coils.

### MukB ATPase activity is stimulated by both *apo*- and *holo*-AcpP

AcpP overexpression in *E. coli* results in a mixture of both *apo-* and *holo*-AcpP species (Supplementary Fig. 4a). Modification of the PPant group through the covalent interaction of acyl groups within the cell, generates a plethora of acylated AcpP intermediates^33^. We therefore investigated whether posttranslational modification of AcpP is required for its interaction with MukB. Analysis of MukB by nMS confirmed the presence of both co-purified *apo-* and *holo*-AcpP within samples (Fig. 1d). Furthermore, we commonly observed additional bands on SDS-PAGE sensitive to reducing agent that ran with a higher molecular mass than purified MukB or the AcpP interacting truncated variants, MukB^HN^ and MukB^N^ (Indicated with an asterisk in Fig. 1c and 2c). Analysis of these bands with anti-AcpP antibody and proteomic MS demonstrated the presence of AcpP (Supplementary Fig. 4b). These bands were also observed in a selection of MukB neck mutants including MukB^G1116E^, MukB^W1117E^ and MukB^V1124E^, but absent in the MukB^C1118E^ sample, suggesting the formation of a disulfide bond between C1118 and the free thiol of *holo*-AcpP (Supplementary Fig. 4b). This disulfide interaction was unnecessary for *in vitro* ATPase stimulation, as both purified *apo*- and *holo*-AcpP could stimulate MukB ATPase to the same extent (Supplementary Fig. 4c). In addition, cells expressing MukB^C1118E^ were viable and displayed apparent WT MukBEF activity (see below). Nevertheless, the formation of this disulfide bond could contribute to the stabilization of the AcpP-MukB interaction.

### MukBEF complexes that are deficient in AcpP binding have perturbed behavior *in vivo*

Next, we assessed the viability of MukB mutants impaired in AcpP binding by transforming plasmid borne copies of the mutants into a Δ*mukB* background strain. Δ*mukB* cells exhibit temperature-sensitive growth in rich medium at 37 °C, which was restored by basal expression from the multi-copy number plasmid pET21a expressing a WT *mukB* gene (Supplementary Fig. 5a). All of the single and double MukB mutants, which were deficient in AcpP binding *in vitro*, had a Muk^+^ phenotype, as assessed by growth at 37 °C. In contrast, cells expressing MukB^KRK^ showed temperature-sensitive growth at 37 °C, consistent with the lack of ATPase activity in this mutant and the substantially impaired response to added AcpP (<25% residual activity in the presence of a 100-fold excess concentration of AcpP; a concentration approaching that *in vivo*) (Fig. 2d). MukB^KR^ (K1114E, R1122E), MukB^RK^ (R1122E, K1125E) and MukB^KC^ (K1114E, C1118E) were Muk^+^, as assessed by growth at 37 °C, indicating that the temperature-sensitivity of MukB^KRK^ is likely due to a lack of AcpP interaction, rather than protein conformational changes induced by the mutations. Consistent with our observations, multiple substitutions in the AcpP-target protein interface are required to abolish AcpP binding with other AcpP binding proteins in addition to MukB^34,35^.

We next explored the functional consequences of the impaired MukB-AcpP interactions by analyzing the behavior of WT and mutant MukBEF complexes by quantitative live cell imaging. We expressed basal levels of MukB and its variants from the multi-copy number plasmid pBAD24 in Δ*mukB* cells containing a functional mYpet fusion to the endogenous *mukE* gene and fluorescent markers located near *oriC* (*ori1*) and close to the centre of *ter* (*ter3*)^15^. In cells expressing WT MukB, fluorescent MukBEF foci were associated with the *ori1* locus, as reported previously by ourselves and others for MukBEF expressed from the endogenous chromosomal locus (Fig. 4a and 4b; 57.1 ± 0.2% colocalization; distances within the diffraction limit (∼264 nm))^3,15,16,36^. Consistent with this, only 7.9 ± 0.3% of MukBEF foci colocalized with *ter3*. In contrast, MukB^EQ^EF foci colocalized with *ter3* and not *ori1*, as reported previously, because they remain associated with MatP-*matS* within *ter*, as a consequence of their defect in ATP hydrolysis^3,15,16^. A MukB mutant that does not bind ATP (MukB^DA^), had its MukBEF distributed over the whole nucleoid, with few, if any, defined fluorescent foci (Fig. 4a)^3,15,16^.

**Figure 4.**
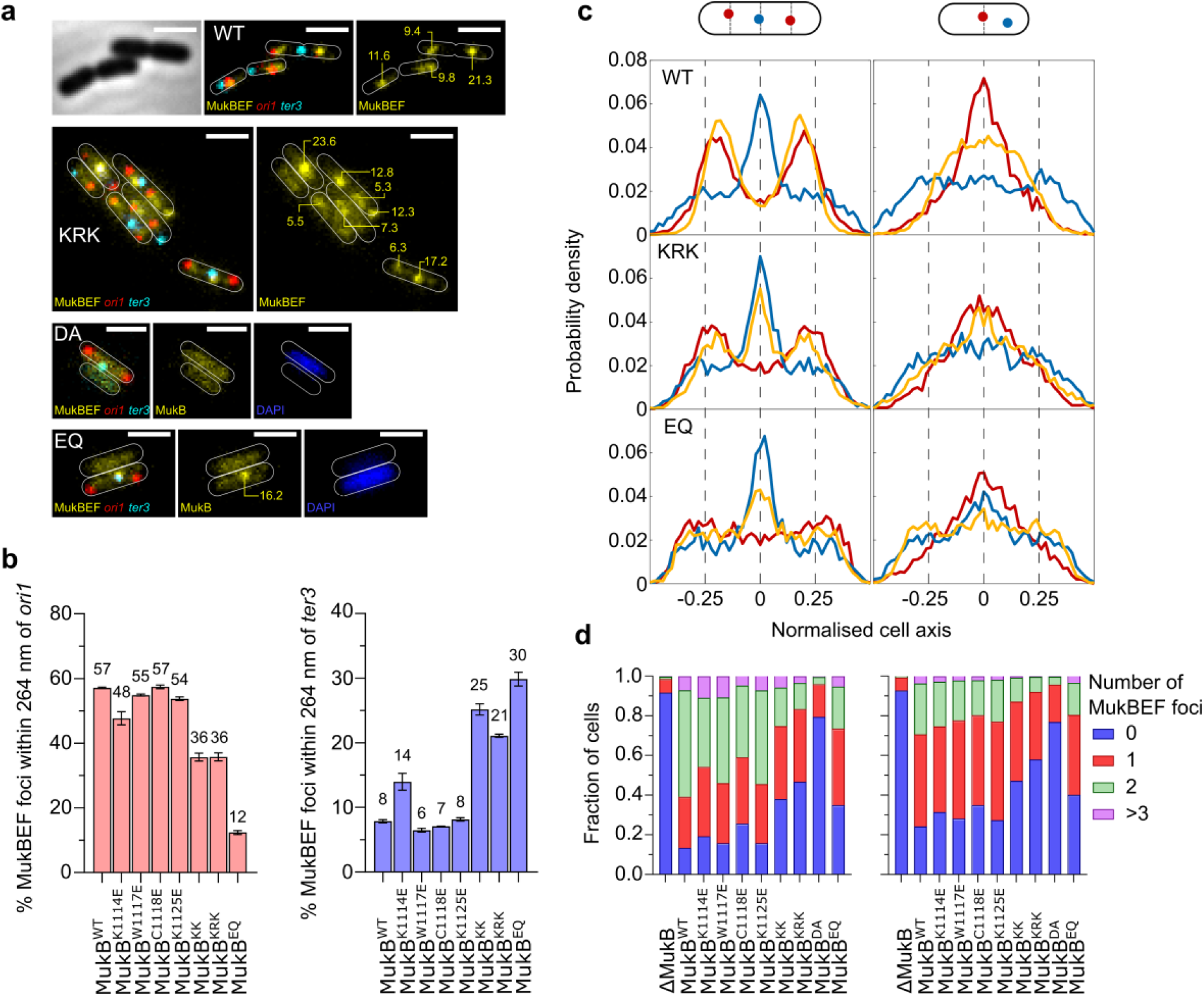
MukBEF complexes that are deficient in AcpP binding have perturbed behavior *in vivo*. Δ*mukB* cells with fluorescently labelled MukE (mYPet), *ori1*(mCherry), and *ter3* (mCerulean) were grown in minimal glycerol medium at 30 °C. Under these conditions, basal expression from a pBAD24 plasmid encoding WT MukB was sufficient to confer a Muk^+^ phenotype on cells. (**a**) Representative images of the indicated strains. The numbers on the images indicate relative brightness of the foci. (**b**) Colocalization of fluorescent MukBEF complexes with *ori1* and *ter3* for the indicated cells (MukB^WT^ 8534 cells, MukB^K1114E^ 7862 cells, MukB^W1117E^ 5402 cells, MukB^C1118E^ 9446 cells, MukB^K1125E^ 9911 cells, MukB^KK^ 5900 cells, MukB^KRK^ 3676 cells and MukB^EQ^ 3849 cells; ± SD from three biological repeats). (**c**) Position of MukBEF foci relative to *ori1* and *ter3*, with respect to the cell axis for all analyzed cells. (**d**) Histograms showing number of fluorescent MukBEF foci/cell with respect to *ori1* and *ter3*. Left panel, cells with 2 *ori*1 loci and 1 *ter3* locus. Right panel, cells with a single *ori1* focus because the locus has not replicated/segregated.

The AcpP binding-impaired variants all produced fluorescent MukBEF foci. They fell into two classes; those indistinguishable from the pattern of WT MukB focus distribution (MukB^W117E^, MukB^C1118E^ and MukB^K1125E^) and those that had a reduced *ori1* association and increased *ter3* association. These latter variants all contained the MukB^K1114E^ mutation either alone, or in combination with one or two further mutations in the AcpP binding region, MukB^KK^ and MukB^KRK^, respectively. MukB^K1114E^, showed a small reduction in association with *ori1* (47.7 ± 2.0 %) and a complementary increase in association with *ter3* (14 ± 1.3%). MukB^KK^ and MukB^KRK^ shared almost identical MukBEF foci properties; 35.7 ± 1.2% and 35.8 ± 1.2% colocalization with *ori1*, respectively, and substantially increased association with *ter3* (25.2 ± 0.9% and 21.0 ± 0.3% t*er3* colocalization, respectively). Despite these similarities only MukB^KRK^ cells exhibited temperature sensitive growth, while the double mutants, like the single ones, grew at 37° C. The behavior of the mutants in relation to *ori1/ter3* localization was independent of whether there was a single *ori1* locus present (in cells soon after birth that had not replicated or segregated the *ori1* locus), or whether there were two sister *ori1* loci, after replication and segregation (Fig. 4c). Nevertheless, we did note that the double and triple mutants had an increasing proportion of cells with no detectable fluorescent MukBEF foci (38 ± 2% and 43 ± 1%, respectively), compared to only 12 ± 2% in WT MukB cells (Fig. 4d), suggesting a significant proportion of cells defective in ATP binding and chromosome association.

The progressive shift from *ori1* to *ter3* co-localization in mutants carrying the MukB^K1114E^ mutation was further evident when the normalized distribution of *ori1, ter3* and MukBEF foci along the longitudinal cell axis was plotted (Fig. 4c). In cells expressing MukB^KRK^ that had 2 *ori1* loci at ¼ and ¾ positions on the long cell axis, a large proportion of MukBEF foci were at the cell center where *ter3* is preferentially located, in addition to the ¼ and ¾ positions. This phenotype is intermediate between cells expressing WT MukB and those expressing MukB^EQ^ (Fig. 4a-c)^15^. The intermediate MukB^KRK^ phenotype was also reflected in a slight shift in *ori1* positioning from the ¼ and ¾ positions towards the poles, which was more evident in MukB^EQ^ cells, as well as in cells lacking MukB (Fig. 4c)^36^. We conclude that MukB^KRK^ cells can still form chromosome-associated MukBEF complexes, but at least a substantial fraction of these are impaired in MukBEF function, consistent with a defect in ATP hydrolysis and consequent preferential location within *ter*.

Cells expressing MukB^G1116E^ and MukB^V1124E^ also exhibited temperature-sensitivity, although the defect was not as complete as for Δ*mukB* cells. <10% of MukB^V1124E^ expressing plated cells yielded colonies at 37 °C, with the surviving colonies being relatively small. A higher proportion of MukB^G1116E^ expressing cells grew at 37 °C, but the colonies were again smaller (Supplementary Fig. 5a). The basis for this sensitivity in MukB^G1116E^ cells is not clear, as cells grown at 30 °C in minimal media had a WT MukB^+^ phenotype as assessed by fluorescent MukBEF foci that are *ori1*-associated and not *ter3*-associated (Supplementary Fig. 5b and c). MukB^G1116E^ expressing cells exhibited a slightly increased fraction of anucleate cells when grown at 30 °C (Supplementary Fig. 5c). In contrast, cells expressing MukB^V1124E^ displayed no clear MukBEF foci, but diffuse mYPet fluorescence similar to cells containing MukB^DA^ (Supplementary Fig. 5b). Despite interacting with AcpP and demonstrating moderate ATPase activity *in vitro*, MukB^V1124E^ seemed unable to interact stably with the chromosome, presumably because the mutation directly interferes with MukB function, consistent with its significant formation of anucleate cells during growth at 30 °C (Supplementary Fig. 5c). The observation that mutations in this region of the MukB coiled-coil can interfere with AcpP binding or otherwise influence MukB function underlines the importance of the joint region in SMC complexes.

## DISCUSSION

We have characterized the specific interaction of AcpP with the joint region of the MukB coiled-coils and have shown that it is necessary for MukB ATPase activity *in vitro* and for normal MukBEF function *in vivo*. The cellular consequences of the MukB-AcpP interaction remain to be determined; in particular, understanding whether AcpP binding to MukBEF *in vivo* is constitutive and unregulated, or whether it is modulated during cycles of MukBEF action, and/or by cellular metabolism. Activation of MukB ATPase activity by AcpP binding, underlines the importance of the joint whose functional roles are only now being revealed. This is emphasized by our demonstration that other mutations in the AcpP binding region of the MukB joint, which do not affect AcpP binding, can perturb MukB function, whether it be impaired ATPase, or *in vivo* action.

The molecular mechanism by which AcpP regulates MukB ATPase activity and overall MukBEF action remains unknown. The AcpP binding site at the MukB joint is relatively distant from the ATPase head and the ‘bent elbow’ configuration of MukB occurs in the absence of bound AcpP^9^. The SMC joint is highly conserved^4,5^ and can be bound by other SMC accessory proteins^37^. Studies of both prokaryote and eukaryote SMC complexes have led to proposals that conformational flexibility in the coiled-coils, facilitated by plasticity of the joint, allows transitions in the disposition of the two SMC heads during their juxtaposition, engagement and disengagement during cycles of ATP binding and hydrolysis. These must be coupled with changes in DNA association during presumed loop extrusion by the complexes^4,5,7^. We favor the view that AcpP binding to the MukB joint modulates such transitions. Since AcpP is acidic and the MukB region involved in its interaction is basic (Supplementary Fig. 1c), it is possible that DNA and AcpP, compete at least transiently, for association with the joint region during these transitions. We have shown that AcpP binding to the MukB joint is not required for MukF binding to MukB, nor is it required for nucleotide- and MukEF-dependent head engagement in the truncated MukB^HN^ variant, as assessed by native gel electrophoresis. Nevertheless, as the disposition of MukB^HN^ ATPase heads are not constrained by the elbow, hinge and the rest of the coiled-coils, this head engagement assay may not reflect the conformational changes that are likely necessary during head juxtaposition and engagement of the full-length protein^4,7,38,39^.

Our observation here that dimer of dimer (DoD) complexes of full length MukB complexed with MukEF, the functional unit *in vivo*^3^, can be detected in the absence of bound AcpP, or AMPPNP-induced head engagement (Fig. 3 and Supplementary Fig. 3), demonstrates that the configuration of two ATPase heads of a MukB dimer prevents two MukF C-terminal domains of a MukF dimer binding to the same MukB dimer, even in the absence of head engagement. Our favored interpretation is that the proximity of the hinge to one of the heads, in the elbow-bent configuration (Fig. 1a), generates an asymmetry, in a way similar to that induced by head engagement^2^, so that only one MukF C-terminal domain can bind a head in a MukB dimer; leaving the other C-terminal domain to capture a second MukB dimer (Fig. 3b). An alternative model in which the disposition of unengaged heads is constrained by relatively rigid coiled-coils in the neck region, again allowing only one MukF C-terminus to bind a MukB dimer, seems less likely.

Given that other SMC complexes can act in the absence of AcpP binding to the joint, it is difficult to rationalize why this requirement has evolved in the MukBEF clade; there is no obvious connection between AcpP and the other MukBEF co-evolved players that include MatP, SeqA, Dam, and topoisomerase IV^15,40^. AcpP is highly abundant (>10^2^-fold cellular molar excess over endogenous MukBEF) and is involved in a wide range of essential steps in fatty acid biosynthesis, along with other specific interactions. Since it exists in a wide range of acylated and unacylated forms, it is challenging to imagine how any modulated MukBEF activity on chromosomes results from cellular changes in AcpP as a consequence of changes in fatty acid metabolism. Parenthetically, MukBEF function only becomes essential for cell viability under condition of rapid growth during which overlapping rounds of replication occur^16^. Indeed, the MukBEF clade of SMC complexes is largely confined to bacteria that support overlapping rounds of replication as part of their lifestyle. Nevertheless, MukBEF is clearly active and important for normal chromosome organization-segregation under conditions of slow growth, when each round of replication is initiated and terminated in the same cell cycle^3,15,16^. Although our work has not identified any specific form of AcpP that preferentially interacts with MukB or influences its activity, any connection between cellular metabolism and the activity of MukBEF complexes on the chromosome, is likely to involve a specific form (or forms) of AcpP whose abundance and activity is under metabolic control. In this scenario, levels of fatty acid biosynthesis could be coordinated in some way with chromosome organization-segregation mediated by MukBEF. Our assays have found no evidence for this; *apo*-AcpP and *holo*-AcpP had comparable activities in stimulating MukB ATPase *in vitro*, while a disulfide between the PPant free thiol and MukB^C1118^ is not essential for either ATPase or *in vivo* function. An alternative scenario to one in which the AcpP-MukB interaction modulates MukBEF action with fatty acid and lipid synthesis is one in which this is an ‘accidental’ recruitment of a protein during evolution, just like the recruitment of the ‘metabolic enzymes’, ArgR, ArcA and PepA, as essential accessory factors in site-specific recombination essential for multicopy plasmid stability^41,42^.

Elsewhere, it has been proposed that the interaction of AcpP with proteins uninvolved in acyl transfer may contribute to the coordination of cellular metabolism. For example, the SpoT-AcpP interaction may help coordinate the cells protein synthesis stringent response to fatty acid starvation^25,28^. Similarly, the interaction between AcpP and the SecA component of the protein membrane translocase machinery could couple fatty acid-lipid metabolism with protein transport through the inner membrane. Although it has been proposed that binding of AcpP to MukB might mediate interactions with the SecA component of the protein membrane translocase machinery, to allow for correct *oriC* positioning within cells^43,44^, in our opinion this appears unlikely. A Turing patterning mechanism positions the largest cluster of MukBEF complexes on the chromosome at either midcell or ¼ positions and the *ori* association with these clusters results directly from the depletion of MukBEF complexes from *ter* as a consequence of their dissociation directed by their interaction with MatP-*matS*^16^. We are unaware of any compelling evidence that replication origins are associated either with SecA complexes or the inner membrane.

The perturbed *ori1* positioning in AcpP binding defective MukB^KRK^ expressing cells is similar to that observed in other situations where MukBEF function is impaired sufficiently to give a temperature sensitive growth phenotype, regardless of whether it is a defect in ATP binding (MukB^DA^), hydrolysis MukB^EQ^), or where there is a complete lack of MukB. The ability of MukB^KRK^ expressing cells to form fluorescent clusters of MukBEF complexes demonstrates that under conditions of impaired AcpP binding, these complexes can still associate with the chromosome, with at least a substantial fraction of these being impaired in MukBEF function, consistent with a defect in ATP hydrolysis and consequent preferential location within *ter*, similar to MukB^EQ^EF complexes that cannot be displaced from *ter* because of their defect in ATP hydrolysis^3,15,16^. Since a proportion of cellular MukB^KRK^ is likely to be bound by AcpP, given the latter’s abundance, we believe this explains why some MukB^KRK^ complexes are *ori*-associated and at least partly functional, albeit with cells having a Muk^-^ phenotype as assessed by temperature sensitivity. In a situation where MukB could not bind AcpP at all, we do not know whether the disposition of the heads would allow sufficient ATP binding to associate with *ter* as in *mukB*^*EQ*^ cells, or whether ATP binding would be so transient that few if any chromosome-associated complexes would be present, as in *mukB*^*DA*^ cells. The work reported here, provides the platform for future studies of the MukBEF mechanism and how it is influenced by AcpP. This will require an integrated combination of structural, biochemical, biophysical and genetic studies and may elucidate more mechanistic and functional insights into the MukBEF clade of proteins, which has evolved an apparently unique architecture, along with a distinctive family of co-evolved partners.

## Methods

### Protein overexpression and purification

MukB-His (and all derivatives thereof), MukE-His and His-MukF were overexpressed from pET vectors and purified as previously described^12^, with the addition of a final step. Following elution from either a HiTrap Heparin HP or HiTrap DEAE FF column (both GE healthcare), appropriate fractions (selected by 4-20% gradient SDS-PAGE) were pooled and concentrated by centrifugal filtration (Vivaspin 20, 5,000 MWCO PES, Sartorius) for loading onto a Superdex 200 Increase 10/300 GL (GE Healthcare) column equilibrated in storage buffer (50 mM HEPES pH 7.3, 300 mM NaCl, 1 mM EDTA, 1 mM DTT and 10% (v/v) glycerol). Peak fractions were assessed for purity (>90%) by SDS-PAGE/Coomassie staining, snap frozen as aliquots and stored at -80 °C.

AcpP was expressed from a pET28a plasmid encoding *acpP* with a thrombin-cleavable N-terminal 6xHis tag in C3031I cells (NEB). 2L cultures of LB supplemented with kanamycin (25 μg/mL) were grown at 37 °C to an OD_600_ of 0.5-0.6 and induced with β-d-1-thiogalactopyranoside (IPTG) at a final concentration of 1 mM. After overnight incubation at 18 °C, cells were harvested by centrifugation, re-suspended in lysis buffer (25 mM HEPES, 150 mM NaCl, 1 mM TCEP, 10 % glycerol) supplemented with a protease inhibitor tablet and homogenized. Cell debris was removed by centrifugation and cell lysate mixed with ∼5 mL of TALON Superflow resin and incubated for 30 mins at 4 °C. The slurry was poured into a column and washed with 10 X volume lysis buffer, 4 X volume wash buffer A (25 mM HEPES, 150 mM NaCl, 1 mM TCEP, 10 % glycerol, 25 mM imidazole) and 1 X volume wash buffer B (25 mM HEPES, 150 mM NaCl, 1 mM TCEP, 10 % glycerol, 100 mM imidazole). Bound proteins were eluted using elution buffer (25 mM HEPES, 150 mM NaCl, 1 mM TCEP, 10 % glycerol, 250 mM imidazole) and dialyzed overnight in lysis buffer with the addition of thrombin protease (10U per 1 mg of AcpP). Uncleaved protein was removed by incubation with TALON Superflow resin before concentrating for loading onto a Superdex 75 Increase 10/300 GL (GE Healthcare) column equilibrated in lysis buffer. Peak fractions were assessed for purity (>90%) by SDS-PAGE/Coomassie staining, snap frozen as aliquots and stored at -80 °C. *P. aeruginosa* AcpH and *B. subtilis* SFP were purified as described^45^.

### Maturation of AcpP

The removal of the AcpP PPant group was achieved as described previously^45^. The addition of the PPant group was achieved in a similar manner, except the final reaction buffer contained 50 mM Tris, pH 7.4, 150 mM NaCl, 10 % glycerol, 0.5 mM TCEP, 1 mM CoA and 0.1 mg/mL *Bs*SFP. After overnight incubation at 37 °C reaction completeness was determined by 20% urea-PAGE. Protein samples were then purified by size exclusion chromatography, snap frozen as aliquots and stored at - 80 °C.

### ATP hydrolysis assays

An EnzCheck Phosphate Assay Kit (ThermoFisher Scientific) was used as described previously^12^, with the exception that all final reactions contained 65 mM NaCl. The reaction was started with the addition of ATP to a final concentration of 1.3 mM.

### Native-state ESI-MS spectrometry

Prior to MS analysis, protein samples were buffer exchanged into 200 mM ammonium acetate pH 8.0, using a Biospin-6 (BioRad) column and introduced directly into the mass spectrometer using gold-coated capillary needles (prepared in-house;). Data were collected on a Q-Exactive UHMR mass spectrometer (ThermoFisher). The instrument parameters were as follows: capillary voltage 1.1 kV, quadrupole selection from 1,000 to 20,000 m/z range, S-lens RF 100%, collisional activation in the HCD cell 50-200 V, trapping gas pressure setting kept at 7.5, temperature 100-200 °C, resolution of the instrument 12500. The noise level was set at 3 rather than the default value of 4.64. No in-source dissociation was applied. Data were analyzed using Xcalibur 4.2 (Thermo Scientific) and UniDec^46^. Data collection for all spectra was repeated at least 3 times.

### Blue-Native gel electrophoresis (BN-PAGE)

MukB or MukB^HN^ (0-4.5 μM) was incubated with MukF (1.5 μM), MukE (3 μM) and AcpP (at the indicated concentrations) in 4X Native PAGE sample buffer (ThermoFisher Scientific, BN2003) with DTT (1 mM) and MgCl_2_ (1 mM) for 30 min at 22 ± 1 °C. Samples were then analyzed using 3-12% native Bis-Tris gels with dark blue cathode buffer. Gels were destained in 40% (v/v) ethanol, 10% (v/v) acetic acid for 30 min before destaining with 8% (v/v) acetic acid overnight.

### Western blot analysis

MukB samples were heated to 95 °C in LDS Sample Buffer (4X) (ThermoFisher NP0007) with or without the presence of reducing agent. Samples were then analyzed using NuPAGE™ 7%, Tris-Acetate SDS-PAGE (ThermoFisher EA03585BOX) followed by western blots using anti-AcpP (LSBio, LS-C370023) as primary and goat anti-rabbit HPR as secondary antibody.

### Proteomics

BS^3^ (50-250X molar excess over MukB^HN^) was added to a sample of MukB^HN^, either co-purified with AcpP or with the addition of recombinant AcpP (at various molar ratios), Reactions were incubated at RT for 30 mins and quenched with Tris buffer (50 mM) before diluting with SDS-loading buffer and analyzed using SDS-PAGE. Protein bands were digested with trypsin overnight at 37 °C as described previously^14^). Peptides were separated by nano-flow reversed-phase liquid chromatography coupled to a Q Exactive Hybrid orbitrap mass spectrometer (Thermo Fisher Scientific). The peptides were trapped onto a C18 PepMap 100 pre-column (inner diameter 300 mm × 5 mm, 100 Ǻ; Thermo Fisher Scientific) using solvent A (0.1% formic acid in water) and separated on a C18 PepMap RSLC column (2 cm, 100 A; Thermo Fisher Scientific) using a linear gradient from 7 to 30% of solvent B (0.1% formic acid in acetonitrile) for 30 min, at a flow rate of 200 ml*/*min. The raw data were acquired on the mass spectrometer in a data-dependent mode. Typical mass spectrometric conditions were: spray voltage of 2.1 kV, capillary temperature of 320 °C. MS spectra were acquired in the orbitrap (m*/*z 350−2000) with a resolution of 70 000 and an automatic gain control (AGC) target at 3 × 10e6 with maximum injection time of 50 ms. After the MS scans, the 20 most intense ions were selected for HCD fragmentation at an AGC target of 50 000 with maximum injection time of 120 ms. Raw data files were processed for protein identification using MaxQuant, version 1.5.0.35 and searched against the UniProt database (taxonomy filter *E. coli*), precursor mass tolerance was set to 20 ppm and MS*/*MS tolerance to 0.05 Da. Peptides were defined to be tryptic with a maximum of two missed cleavage sites. Protein and peptide spectral match false discovery rate was set at 0.01.

### Functional analysis *in vivo*

The ability of MukB variants to complement the temperature-sensitive growth defect of a *ΔmukB* strain was tested as described previously, using basal levels of MukB expression from plasmid pBAD24^12^. Live-cell imaging used cells grown in M9 minimal medium with 0.2% (v/v) glycerol, 2 μg ml-1 thiamine, and required amino acids (threonine, leucine, proline, histidine and arginine; 0.1 mg ml-1) at 30 °C. An overnight culture was diluted ∼1000-fold and grown to A_600_ 0.05–0.2 and deposited on a medium containing agarose pad after staining with 1 μg/mL DAPI. The Δ*mukB* cells used had a functional mYpet fusion to the endogenous *mukE* gene, fluorescently labelled *ori1* (mCherry), and *ter3* (mCerulean) (AU2118; *lacO240* @*ori1* (3908) (*hyg*), *tetO240*@*ter3* (1644) (*gen*), Δ*leuB::Plac-lacI-mCherry-frt*, Δ*galK::Plac-tetR-mCerulean-frt*, Δ*araBAD* (AraC+), *mukE*-*mYPet-frt*-*T1-T2*-*Para*-*ΔmukB*-*kan*)^15,16^, expressing basal levels of pBAD24 plasmid-borne WT MukB, the indicated MukB mutants, or empty pBAD24 plasmid control (Δ*mukB)*. Epifluorescence images were acquired on a Nikon Ti-E inverted microscope equipped with a perfect focus system, a 100× NA 1.4 oil immersion objective (Nikon), an sCMOS camera (Hamamatsu Flash 4), a motorized stage (Nikon), an LED excitation source (Lumencor SpectraX) and a temperature chamber (Okolabs). Fluorescence images were collected with 100 ms exposure time using excitation from a LED source. Phase contrast images were collected for cell segmentation. Images were acquired using NIS-Elements software (Nikon). Cell segmentation and spot detection from the fluorescence channel were performed using SuperSegger^47^. Low quality spots were filtered out with a fixed threshold for all data sets (4.5). The threshold was selected to minimise the number of falsely identified MukBEF foci within background signal yet maximise the number of foci analyzed; the threshold ensured ∼90% of cells expressing WT MukB contained at least one MukBEF focus, whilst ∼90% of Δ*mukB* cells had none. The percentages of cells containing one or more spots, distances to the closest *ori1*/*ter3* marker, and localisation along the long cell axis were calculated using MATLAB (MathWorks) as described^16^. For anucleate cell percentages, cells deemed anucleate by DAPI staining and lack of *ori1* marker were counted manually.

## DATA AVAILABILITY

All digital forms of the data are available on request. All materials and analysis codes are available upon reasonable request.

## AUTHOR CONTRIBUTIONS

J. P.P., L.K.A. and D.J.S. conceived and directed the project. J. P.P, G. L.M.F. and J.R.B. undertook biochemical experiments. J. M. helped with quantitative imaging analysis. C.V.R. provided facilities for nMS. The paper was drafted by J. P.P, L.K.A. and D.J.S., with all authors participating in the final manuscript.

## ACKNOWLEDGEMENTS

We thank all members of the Sherratt lab. for useful discussions, the departmental proteomics unit for proteomics support and Rachel Baker for excellent technical support. We thank Frank Bürmann, Jan Löwe (MRC LMB, Cambridge, UK) and Mike Burkart (UCSD, San Diego) for helpful discussions.

## Supplementary Material

**Supplementary Figure 1.**
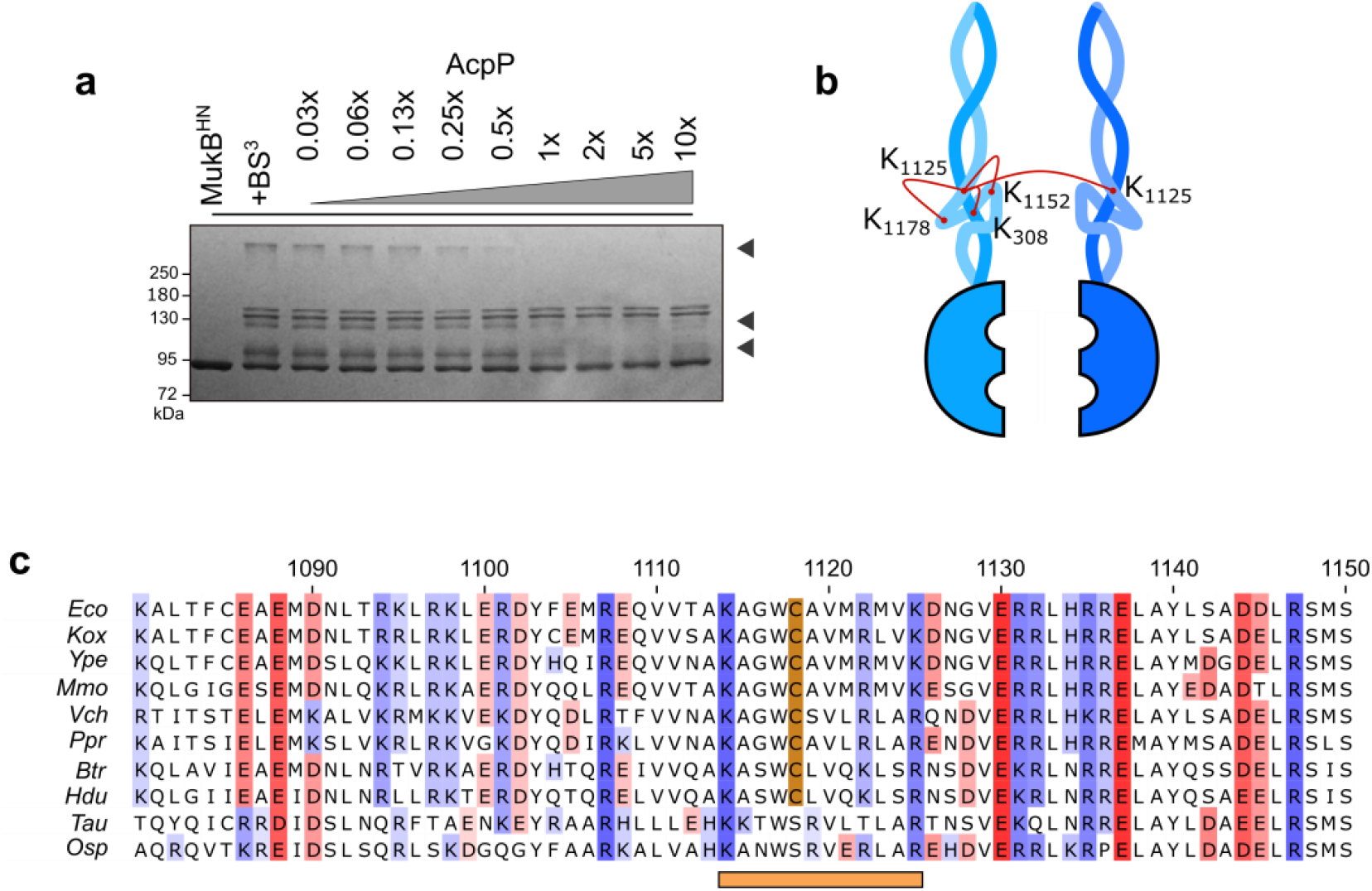
Characterization of the MukB-AcpP interaction. (**a**) Identification of BS^3^ cross-links, both inter- and intra-molecular cross-linking of MukB^HN^ is inhibited by AcpP (indicated by grey arrows). (**b**) Schematic of the cross-links in MukB^HN^ that are inhibited by AcpP binding. (**c**) Alignment of MukB sequences in the region bound by AcpP. Conservation in acidic and basic residues are indicated in red or blue respectively. The horizontal bar indicates the region analyzed in the work here. The conserved C1118 residue is also highlighted. Eco – *Escherichia coli*, Kox – *Klebsiella Oxytoca*, Ype – *Yersinia pestis*, Mmo – *Morganella morganii*, Vch – *Vibrio cholerae*, Ppr – *Photobacterium profundum*, Btr – *Bibersteinia trehalosi*, Hdu *– Haemophilus ducreyi*, Tau – *Tolumonas auensis*, Osp – *Oceanimonas sp*.

**Supplementary Figure 2.**
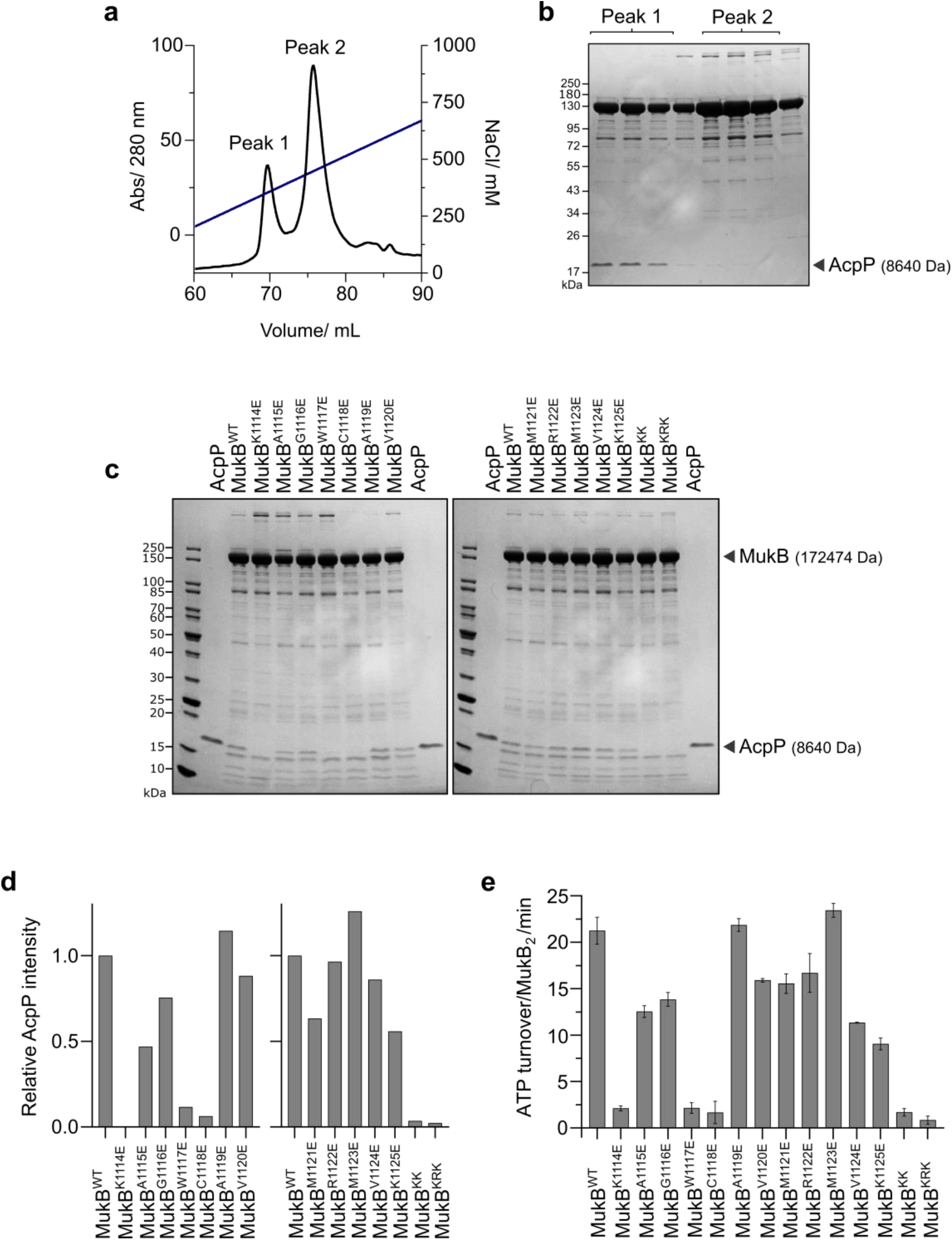
Mutagenesis in the MukB-AcpP interface hinders AcpP co-purification and results in reduced ATPase activity. (**a**) Typical separation of MukB (peak 2) from MukB-AcpP (peak 1) using a salt gradient on a heparin column. (**b**) SDS-PAGE analysis of the peaks in (**a**). (**c**) SDS-PAGE analysis of TALON-purified MukB proteins, indicating the presence or absence of copurified AcpP. (**d**) Relative levels of AcpP associated with the indicated MukB proteins in (**c**), and (**e**) their ATPase levels (±SD from 3 technical repeats).

**Supplementary Figure 3.**
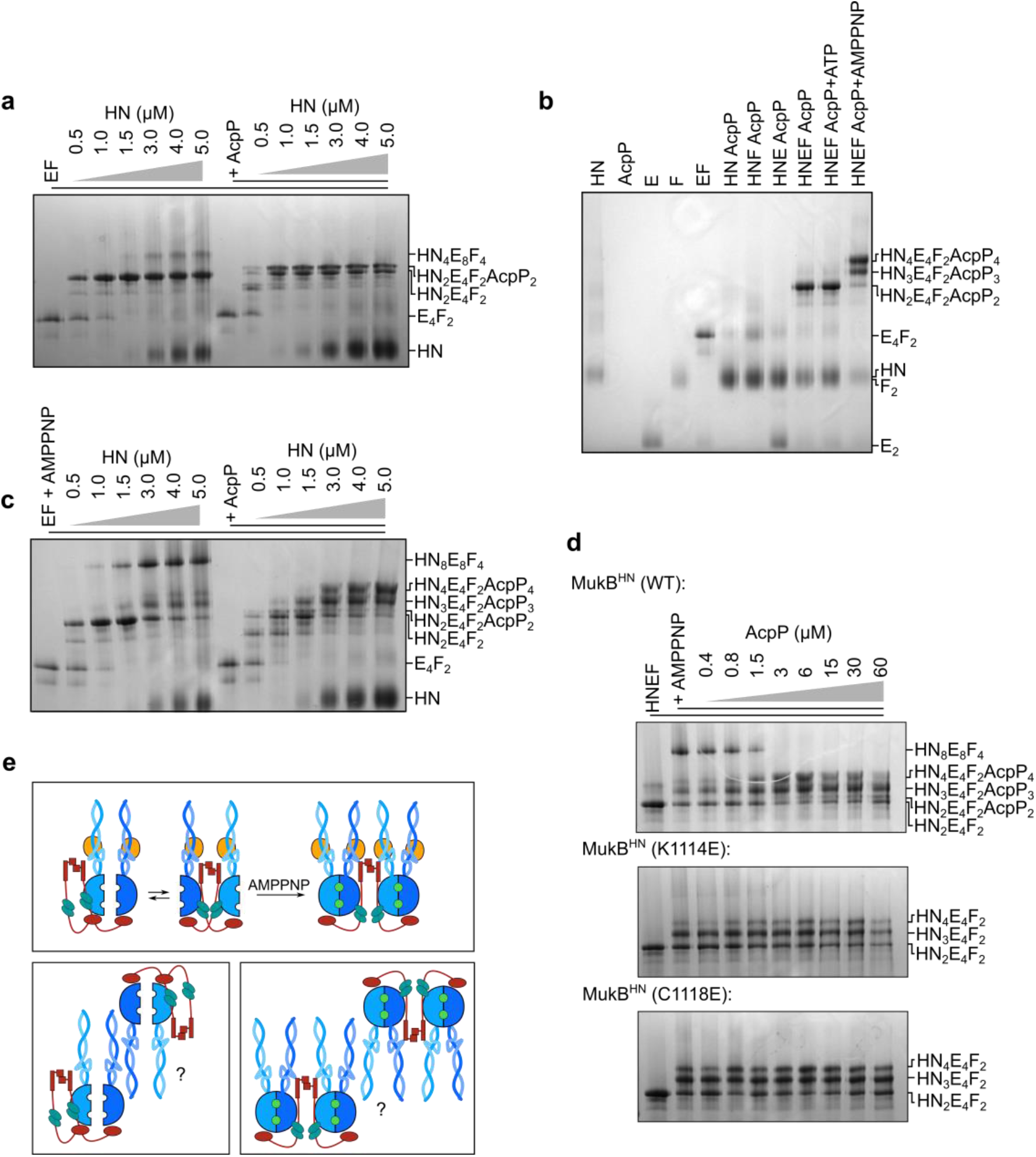
AcpP prevents a coiled-coil interaction in MukB^HN^. BN-PAGE analysis of complex formation in MukB^HN^EF-AcpP. (**a)** AcpP hinders the formation of higher order MukB^HN^EF complexes. (**b**) AMPPNP induces head engagement to form MukB^HN^_3/4_E_4_F_2_ complexes. (**c**) AcpP hinders the formation of AMPPNP-dependent higher order MukB^HN^EF complexes (**d**) AcpP or mutagenesis in the MukB-AcpP interface hinders the formation of higher order complexes. (**e**) Schematic of nucleotide induced MukB^HN^EF-AcpP head engagement (Top), or possible higher order complexes formed through coil-coil interactions in the absence of AcpP in head unengaged (Bottom, left) and head engaged (Bottom, right) complexes.

**Supplementary Figure 4.**
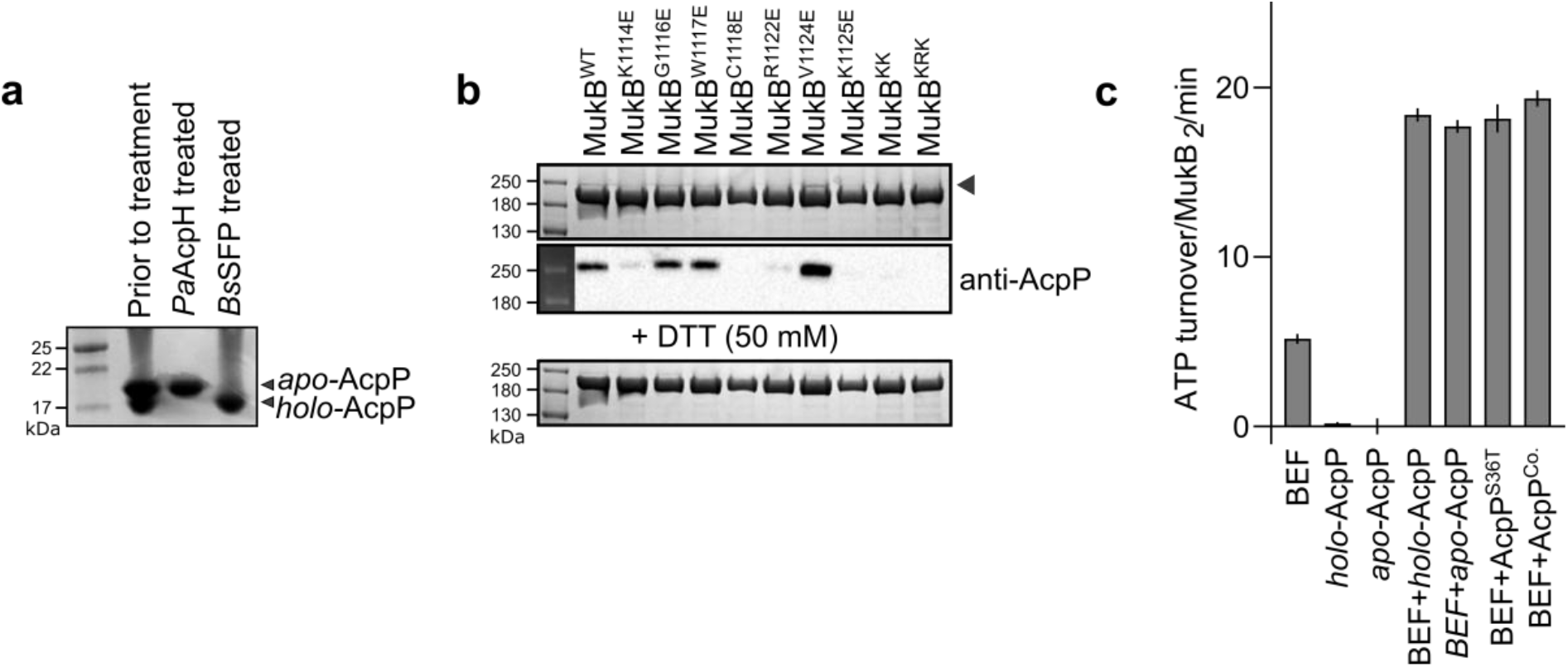
Activities of *apo*-AcpP and *holo*-AcpP. (**a**) 20% urea-PAGE analysis of recombinant AcpP. Overexpression results in the production of *apo-* and *holo*-AcpP. (**b**) SDS-PAGE and western blot analysis of putative disulfide linked MukB-AcpP complexes (indicated by an arrow). (**c**) Initial ATPase activity measurements of MukB in the presence of various AcpP species (±SD from 3 technical repeats). For these experiments, AcpP that still contained the 6XHis-tag was used as its presence has no impact in the observed MukB ATPase activity.

**Supplementary Figure 5.**
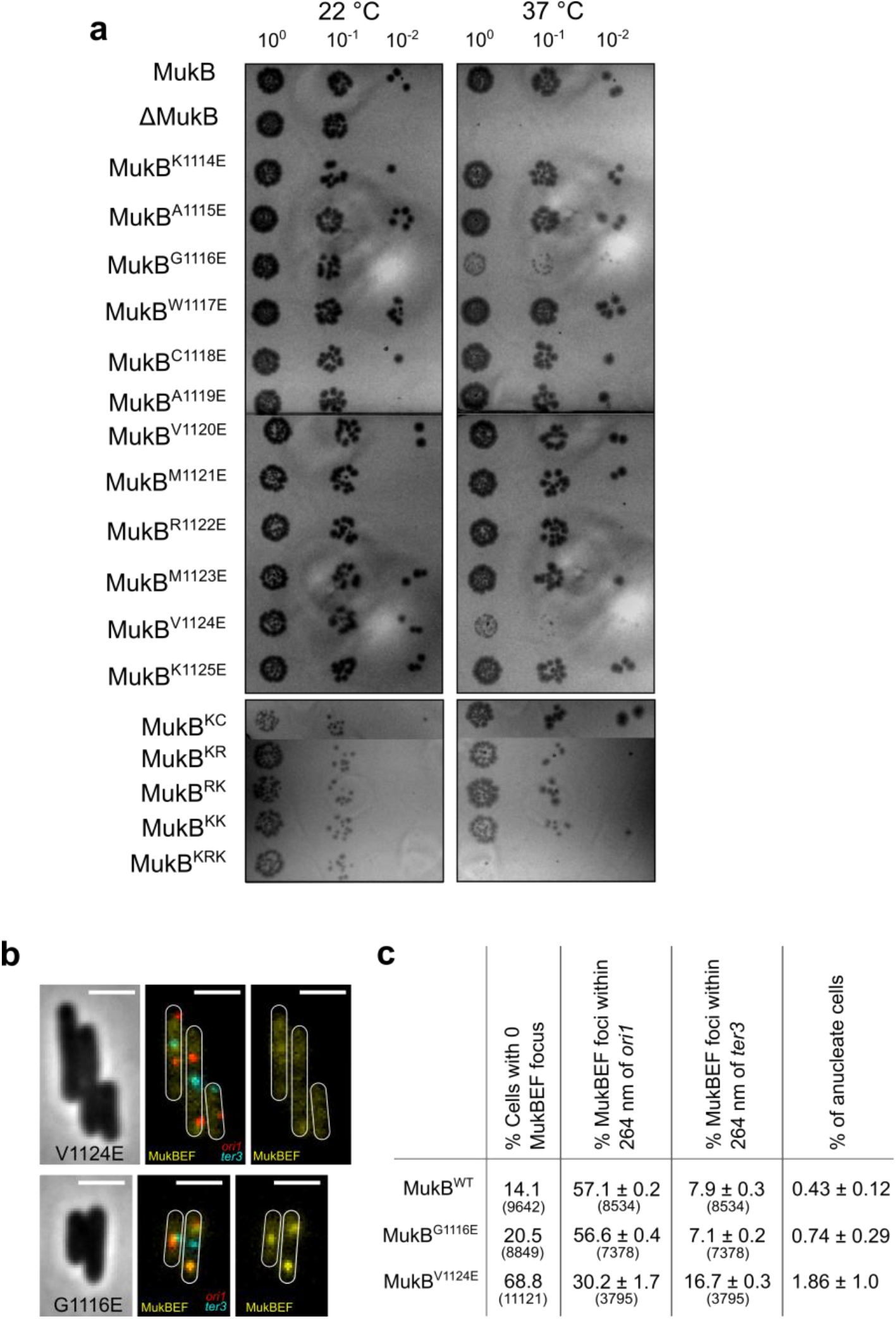
Phenotypes of cells expressing MukB^V1124E^ and MukB^G1116E^ mutants. (**a**)Temperature sensitivity of WT and MukB mutant strains. Cells were grown in LB at 22 °C overnight, diluted and 20 μL spots of the dilutions plated and incubated as indicated. (**b**) Representative images of cells expressing MukB^V1124E^ and MukB^G1116E^ mutants (conditions as in Figure 4). (**c**) Analysis (as in Figure 4) of MukBEF foci in relation to *ori1* and *ter3* loci, and frequency of anucleate cells (± SD from 3 biological repeats. Number of cells analyzed in parentheses.

